# Similarities Between Antibiotic-Resistant *Escherichia coli* from Raw Meat, Commercial Raw Dog Food and Those Causing Extraintestinal Human Infections: A Contemporaneous Geographically Focussed Genomic Epidemiology Study

**DOI:** 10.1101/2024.03.03.583175

**Authors:** Jordan E. Sealey, Beth Astley, Katie L. Sealey, Aimee M. Daum, Defne Kiziltan, Laura Wright, Philip Williams, Matthew B. Avison

**Affiliations:** School of Cellular & Molecular Medicine, University of Bristol. UK; University Hospitals Bristol and Weston NHS Foundation Trust, Bristol. UK

## Abstract

**Objectives:** There is strong evidence linking raw meat or raw dog food (RDF) feeding with dogs shedding antibiotic-resistant (ABR) *Escherichia coli*. As well as potential risks of ingesting ABR *E. coli* when handling/consuming meat, therefore, people who raw-feed pets may face additional risks. However, phylogenetic evidence closely linking *E. coli* found on raw meat/RDF with those causing opportunistic human infections is lacking, partly due to non-contemporaneous sampling. Here, we monitored ABR *E. coli* in meat/RDF in Bristol, UK in 2022/23, and compared *E. coli* isolates with *E. coli* causing human urinary tract and bloodstream infections in Bristol in 2023.

**Methods:** We monitored ABR *E. coli* positivity in raw chicken, beef, pork and lamb meat from 15 large-chain stores and chicken-based RDF from 15 pet stores and selected 200 ABR *E. coli* for Illumina WGS. *E. coli* causing 1182 human infections were sequenced, irrespective of resistance phenotype.

**Results:** Chicken/RDF were most likely to carry amoxicillin-, spectinomycin-, streptomycin-, cefotaxime-, and ciprofloxacin-resistant *E. coli*. Phylogroup A/B1 and B2 *E. coli* dominated meat/RDF and human isolates, respectively. Resistance to antibiotics used predominantly in farmed animals dominated meat/RDF isolates; resistance to antibiotics extensively used in humans dominated human isolates. Nineteen meat/RDF-infection *E. coli* pairs differed by <50 SNPs across the core-genome, with four pairs (three ST69 and one ST117) differing by <20 SNPs and carrying identical resistance genes, suggestive of recent sharing.

**Conclusions:** Using contemporaneous, geographically focussed sampling, we have confirmed close phylogenetic relationships between meat/RDF and human infection-causing ABR *E. coli*. Despite representing a small number of infections, these findings support the consideration of tighter microbiological standards for RDF given that it is explicitly sold to be fed uncooked. Improved risk communication to pet owners and the inclusion of RDF within integrated ABR surveillance is warranted.

## Introduction

*Escherichia coli* is an almost ubiquitous commensal in the gastrointestinal tracts of humans and other warm-blooded animals, and an important pathogen (1). Enteropathogenic *E. coli* causes food-borne diarrhea outbreaks, while extraintestinal pathogenic *E. coli* are leading causes of urinary tract infection (UTI) and bloodstream infection (BSI); these infections are opportunistic and predominantly arise from commensal gastrointestinal flora (2). Antibiotic resistant (ABR) *E. coli*, therefore, pose a significant health threat and are an important indicator for One Health ABR transmission (3).

Meat can become contaminated with fecal bacteria, including *E. coli*, during slaughter and processing (4). In the UK, regulations ensure that raw meat sold for consumption by people and domestic pets meet the same bacteriological limits. For commensal *E. coli*, acceptable limits may reach hundreds of bacteria per gram. Therefore, poor hygiene when handling, storing and preparing raw meat as well as consuming undercooked meat can lead to the ingestion of *E. coli*, including ABR strains (5).

Raw meat feeding of pets has increased in popularity, largely driven by owners’ perceptions of enhanced pet wellbeing (6). A recent UK survey found that around 50% of owners who raw feed their dogs feed commercial raw dog food (RDF) and 50% prepare it themselves, using raw meat sold for human consumption (6). Like raw meat, RDF is frequently contaminated with ABR *E. coli*, including those resistant to the critically important third-generation cephalosporins (3GCs) or fluoroquinolones (FQs) (7-12).

We have demonstrated that dogs who are fed raw meat in the city of Bristol, UK, are more likely to excrete 3GC-resistant (3GC-R) or FQ-resistant (FQ-R) *E. coli* (13,14). Other studies performed in other geographical regions support our findings (15-18). Therefore, people feeding raw meat face multiple ABR *E. coli* exposure risks: handling meat contaminated with ABR *E. coli* and interacting with dogs which are more likely to excrete them (7). However, there remains a scarcity of high-resolution, contemporaneous, geographically focused studies that phylogenetically compare ABR *E. coli* from raw meat and human clinical infections, with none involving RDF. Here we compared 200 ABR *E. coli* from raw meat purchased from large-chain grocery stores and commercial RDF sold in the city of Bristol, UK in 2022-23 with 1,182 *E. coli* isolates from human UTI and BSI in Bristol in 2023.

## Materials and Methods

### Sample collection

We sampled five locations, one per week during October and November 2022 (approximately 25,000 population in each location), within Bristol, a city in South West England. For each location, three large chain grocery stores were selected, and one packet each of raw minced beef, minced lamb, chicken thighs and pork chops were purchased at each store. In total, fifteen samples per meat type were collected, except pork (13 samples due to availability issues). Samples were kept at 4°C and processed within the “use by” date.

Five Bristol-based RDF stores were also identified; locations overlapped with the sampled large-chain grocery stores. Three brands of chicken-based RDF were purchased in each store during September 2023, totaling 15 RDF samples across 11 brands. As RDF is sold frozen, samples were defrosted at 4°C on the day of purchase and processed the next day.

### Microbiology

Under sterile conditions, 200 g meat was bagged (Stomacher^®^ 400 Classic Standard) alongside 200 mL of *E. coli* Enrichment Broth (Oxoid LTD). Sealed bags were placed into screw-top plastic jars for incubation for 5 h at 37°C and 180 rpm. Enriched culture (500 µL) was mixed with glycerol (25% v/v final), twenty microliters of each mixture was spread onto Tryptone Bile X-Glucuronide agar without antibiotic, or with amoxicillin (8 mg/L), spectinomycin (32 mg/L), streptomycin (32 mg/L), amoxicillin/clavulanate (8/2 mg/L), cefotaxime (2 mg/L), and ciprofloxacin (0.5 mg/L). Culture pates were incubated at 37°C for 18 h. Up to 10 blue colonies per plate were tested against the same antibiotics to determine resistance phenotype. One isolate per resistance phenotype per sample was selected for whole genome sequencing (WGS).

### Human infection isolates

Isolates were from blood or urine samples deduplicated by patient. They were collected in 2023 at Severn Pathology, a diagnostic laboratory within Bristol. BSI isolates (n=795) were collected throughout 2023 from hospitalised patients. Community UTI isolates (n=387) were collected for one week in December 2023. All isolates were sequenced, irrespective of antibiotic susceptibility profile.

### WGS and phylogenetic analysis

WGS was performed by MicrobesNG to achieve a minimum 30-fold coverage using standard protocols (https://www.microbesng.com/). WGS used an Illumina NovaSeq 6000 (Illumina, San Diego, USA) with a 250 bp paired end protocol. Data were analyzed using ResFinder 4.7.2 (19), and MLST 2.0.9 (20). WGS data were excluded due to quality control issues if >500 contigs, <50 or >51% GC or <4.5 million or >6 million nucleotides total length. Multiple genomes with the same sequence type (ST) and antibiotic resistance gene (ARG) complement from a single meat/RDF sample were deduplicated.

Phylogenetic analysis used the Bioconda environment. Alignments (**Table S1** shows references) used Snippy and Snippy-Core (https://github.com/tseemann/snippy); SNP distances were determined using SNP-dists (https://github.com/tseemann/snp-dists). Maximum likelihood trees were produced using RAxML with the GTRGAMMA model of rate of heterogeneity (21) and illustrated using Microreact (22).

## Results

### Sample-level positivity for ABR E. coli in raw meat

Eighty-one percent of meat from large-chain grocery stores tested positive for *E. coli*. Positivity was highest in chicken (100%) and almost all chicken samples contained streptomycin-, spectinomycin- and amoxicillin-resistant *E. coli*, significantly greater (Fisher’s Exact *p*<0.05) than that found in beef, lamb or pork. Notably, 47% of chicken samples carried FQ-R *E. coli*, significantly higher than lamb and pork, but not beef. Seven percent of chicken and 13% of beef samples contained 3GC-R *E. coli*; no lamb or pork samples were positive (**Tables 1, S2**).

**Table 1.**
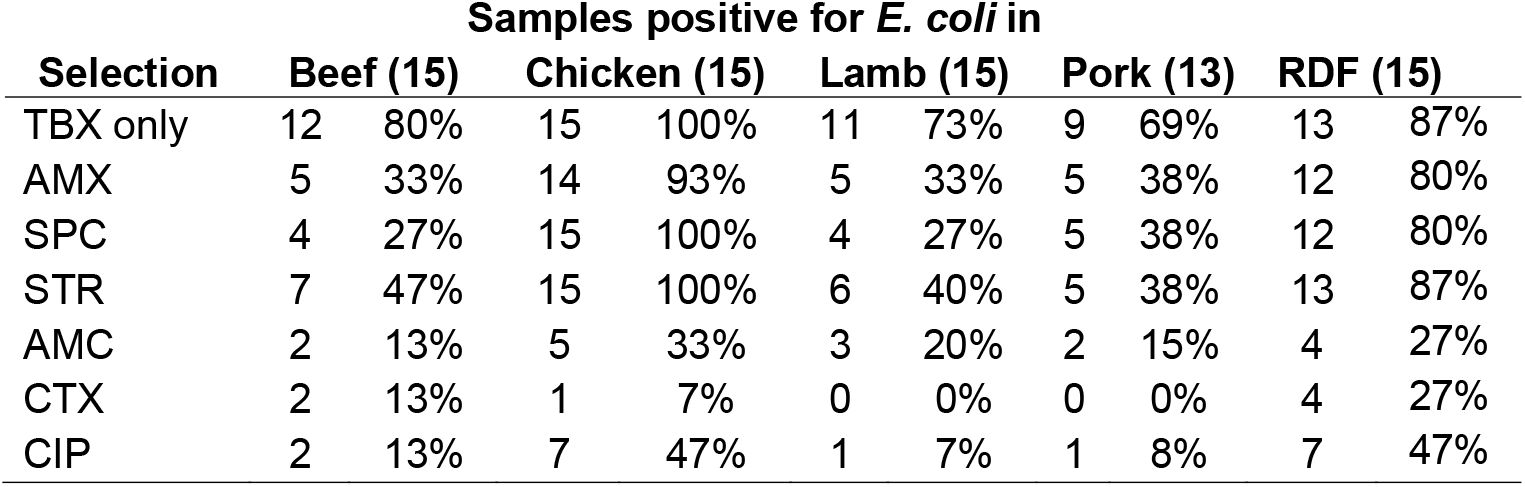
Sample-level positivity for *E. coli* in meat sold primarily for human consumption at large chain grocery stores and in chicken-based RDF sold at specialist pet stores. Total samples tested in brackets.

| Selection | Samples positive for *E. coli* in |  |  |  |  |  |  |  |  |  |
| --- | --- | --- | --- | --- | --- | --- | --- | --- | --- | --- |
|  | Beef (15) |  | Chicken (15) |  | Lamb (15) |  | Pork (13) |  | RDF (15) |  |
| TBX only | 12 | 80% | 15 | 100% | 11 | 73% | 9 | 69% | 13 | 87% |
| AMX | 5 | 33% | 14 | 93% | 5 | 33% | 5 | 38% | 12 | 80% |
| SPC | 4 | 27% | 15 | 100% | 4 | 27% | 5 | 38% | 12 | 80% |
| STR | 7 | 47% | 15 | 100% | 6 | 40% | 5 | 38% | 13 | 87% |
| AMC | 2 | 13% | 5 | 33% | 3 | 20% | 2 | 15% | 4 | 27% |
| CTX | 2 | 13% | 1 | 7% | 0 | 0% | 0 | 0% | 4 | 27% |
| CIP | 2 | 13% | 7 | 47% | 1 | 7% | 1 | 8% | 7 | 47% |

We found no statistically significant difference in ABR *E. coli* positivity among chicken-based RDF samples, compared with grocery store chicken (Fisher’s Exact *p*>0.05; **Tables 1, S2, S3**).

### *Molecular epidemiology of ABR* E. coli from raw meat

In total, WGS for 200 unique ABR *E. coli* isolates were retained: 90 from RDF, 66 (chicken), 17 (beef), 13 (lamb), and 14 (pork). A wide range of *E. coli* STs were detected (**Table S4**). ST10, ST162, ST69, ST88, ST117 and ST1718 were found in multiple meat types, but none in all types. Isolates from ST131, a common human extraintestinal pathogenic ST, were found in one beef (serotype O16:H5 *fimH41*) and one pork (O15:H4 *fimH22*) sample. Chicken-based RDF shared 13 STs with chicken meat, with a statistically significant ST clustering between chicken versus chicken-based RDF compared with all other meat sources combined (Fisher’s Exact *p*<0.001) (**Table S4**).

Following a core genome alignment, a difference of <20 SNPs between two core genomes was used to define clonality, suggestive of recent transmission (23). Thirty clones were identified of which 15 included isolates from more than one meat type (**Table S5**). Considering all pairwise core genome alignments, pork and lamb were significantly less likely to share clonal isolates with chicken-based RDF than chicken (Fisher’s Exact *p*=0.004) and beef (*p*=0.0001), with chicken and beef not being significantly different in this regard (*p*=0.11). Though chicken/RDF sharing was seen in 12 clones and beef/RDF sharing in only five (**Table S5**).

Across the 15 RDF samples, three were from the same brand, and across seven STs we detected multiple isolate pairs <20 SNPs apart between these three samples. Two further brands were each sampled twice, although no <20 SNP links were observed. The remaining eight brands were each represented by a single sample.

### Comparison of ABR E. coli from raw meat sold in Bristol with E. coli causing human infections in the same region

Twenty-one STs were shared between meat and human infection isolates with 47/90, 33/60 and 20/44 isolates, respectively from RDF, chicken and red meat (beef, pork and lamb combined) being of STs that are also seen among human clinical isolates (**Table S6**). These proportions were not significantly different based on type of meat (Fisher’s Exact *p*>0.05).

When considering *E. coli* phylogroups (**Table S7**), phylogroup B2 was significantly more common among human infection isolates than among isolates from poultry (chicken or RDF) or red meat (Fisher’s Exact *p*<0.00001 for each). In contrast, phylogroups A/B1 isolates were significantly less abundant in human isolates compared with poultry or red meat (Fisher’s Exact *p*<0.00001 for each). Phylogroup D was also significantly less abundant in human isolates than in poultry isolates (Fisher’s Exact *p*=0.012) but the difference (13.8% versus 22.4%) was much less than for A/B1 isolates (13.5% versus 66.0%).

We identified 19 incidences where *E. coli* from meat were <50 SNPs different from a paired isolate causing a human infection, with four of these meat/infection pairs being <20 SNPs apart (**Table S8**). Overlaps between meat and human infection isolates from ST162, ST88 and ST10 are illustrated in **Figures S1-S3** with STs ST69 and ST117 – having three and one <20 SNP distanced pairs, respectively – being illustrated in **Figures 1 and 2**.

**Figure 1.**
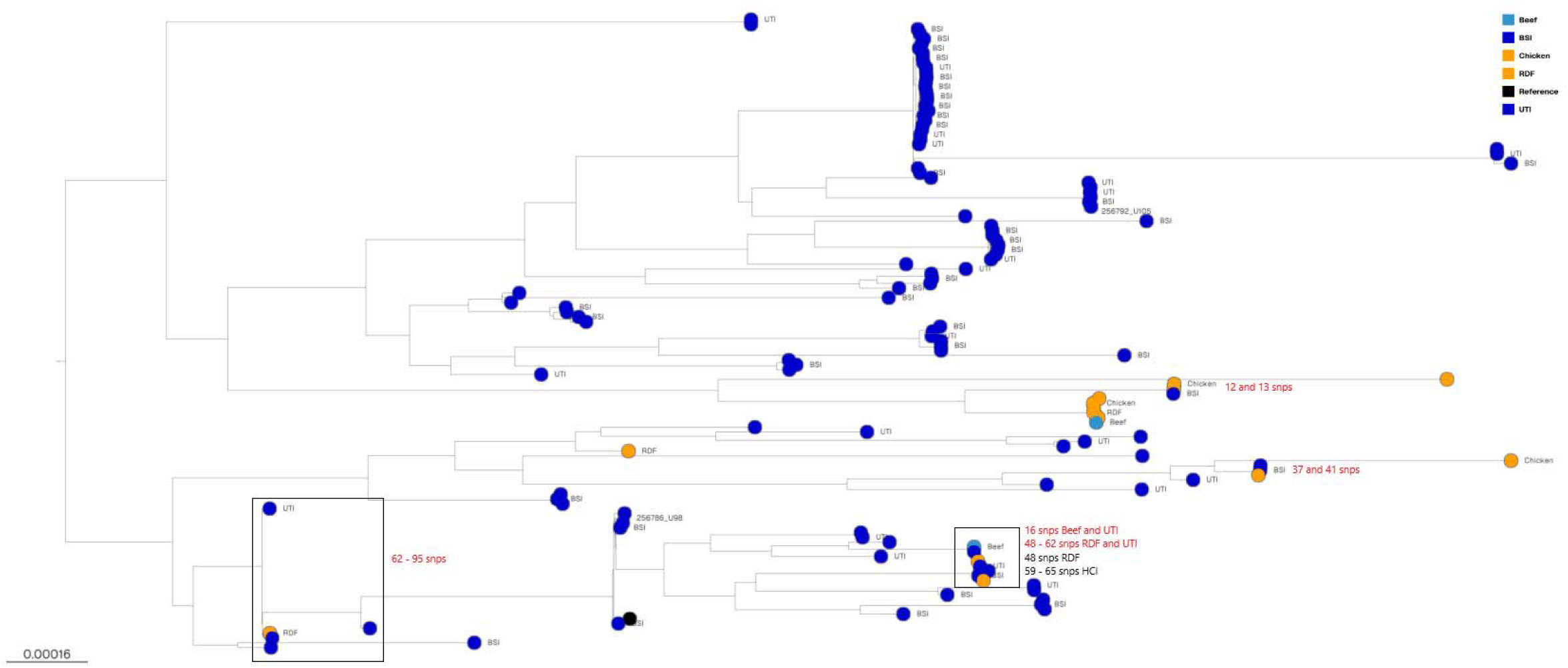
Mid-rooted phylogenetic tree of *E. coli* ST69 isolated from raw meat products (beef (blue), poultry (chicken and RDF, yellow)) and from human clinical infection (HCI; bloodstream and urinary tract infections, blue). Single nucleotide polymorphisms <150 SNPs in red are SNPs between raw meat source and human source, SNPs between isolates from same source are in black.

**Figure 2.**
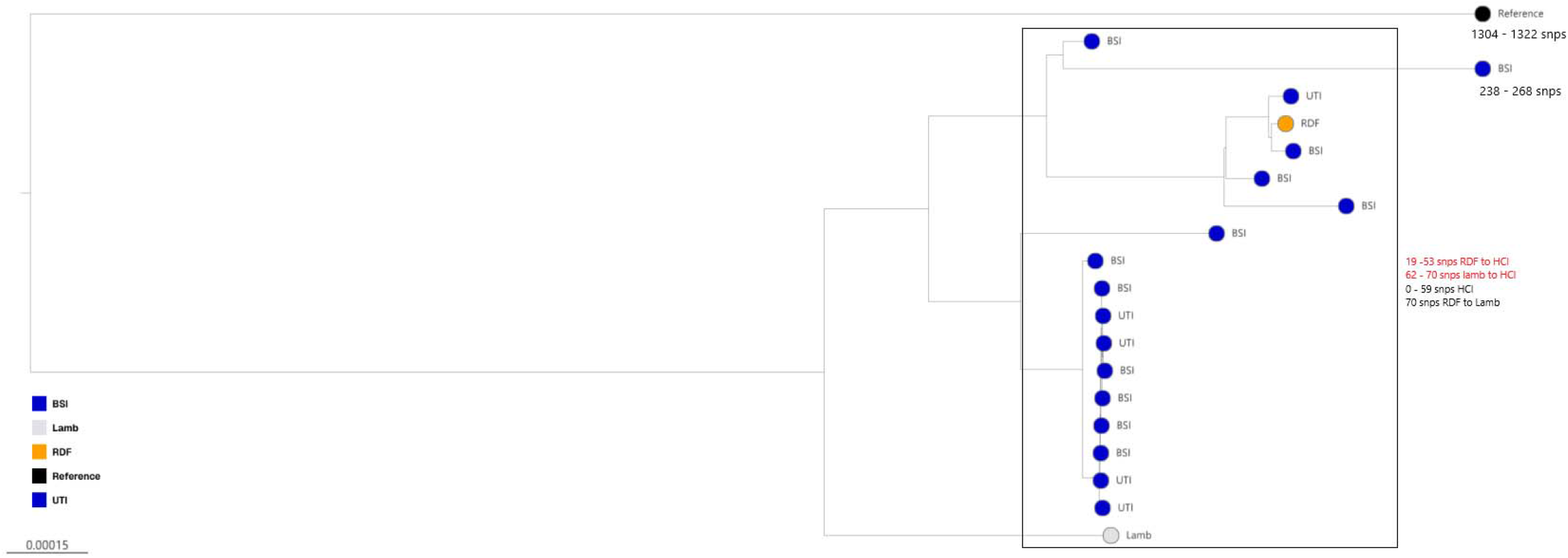
Mid-rooted phylogenetic tree of *E. coli* ST117 isolates from human clinical infections (BSI or UTI, blue), one from raw dog food (RDF, yellow) and another from lamb (grey). Single nucleotide polymorphisms in red are SNPs between raw meat source and human source, SNPs between isolates from one source in black.

### ARG Characterization of Resistant E. coli from Meat and Human Infections

The average number of ARGs per isolate was 5.5 in RDF, 4.7 in chicken, 5.2 in beef, 5.0 in pork, and 3.2 in lamb. Common ARGs detected across most meat sources included *aadA, aph(3’’)-Ib and aph(6)-Id (strAB), sul1/2, dfrA, bla*_TEM-1_, *tetA/B* and *qnrS1* (**Table S9**). It was very rare to find meat isolates within one of the 30 clones defined above having identical ARG complements (**Table S5**). However, the two chicken meat/BSI ST69 pairs differing by 12 and 13 SNPs, respectively all carried *aac(3)-VIa, aadA1, sul1, sul2* and the beef meat/UTI ST69 pair differing by 16 SNPs both carried *strAB, bla*_TEM-1_, *sul1, sul2, tetA, dfrA7*.

FQ resistance was predominately due to chromosomal mutations in meat isolates, with the most common being the triple combination of *gyrA*D87N, *gyrA*S83L *and parC*S80I. ARGs conferring 3GC resistance, were *bla*_CTX-M-1_, *bla*_CTX-M-15_, and *bla*_SHV-12_ with one being a chromosomal *ampC* hyperproducer (**Table S9**).

In the UK, many antibiotics used in livestock are also used in human medicine, e.g. beta-lactams, trimethoprim, sulfonamides, gentamicin, tobramycin and FQs (though the last three are used very rarely in livestock). However, other antibiotics, e.g. florfenicol, neomycin, streptomycin, spectinomycin, and tetracycline are widely used in livestock while use in human medicine is low (though tetracycline derivatives are widely used). By comparing the number of ARGs relevant for each of these main groups of antibiotics in *E. coli* isolates from poultry (chicken and RDF) or red meat (beef, lamb and pork) with those from human infections, we noted that for the antibiotics widely used in human medicine there was either no significant difference in abundance between meat and human infections, or there was a significant overabundance in human isolates. However, for antibiotics predominantly used to treat farmed animals, in most cases there was a significant relative overabundance in isolates from meat (**Table 2**).

**Table 2.** ARGs from meat and human sources conferring resistance to different antibiotic classes.

| Antibiotic | Poultry | Red Meat | Human | Human versus Poultry | Human versus Red Meat |
| --- | --- | --- | --- | --- | --- |
| Beta-lactam | 139 | 37 | 588 | 0.038 | NS |
| Fluoroquinolone | 17 | 3 | 76 | NS | NS |
| Gentamicin/<br>Tobramycin | 19 | 2 | 90 | NS | NS |
| Trimethoprim | 71 | 13 | 353 | 0.006 | 0.01 |
| Sulphonamide | 130 | 30 | 482 | NS | NS |
| Tetracycline | 88 | 33 | 290 | NS | 0.007 |
| Florfenicol | 41 | 6 | 31 | <0.00001 | 0.019 |
| Neomycin/<br>Spectinomycin/<br>Streptomycin | 241 | 66 | 744 | 0.023 | 0.048 |
| Total | 746 | 190 | 2654 |  |  |

Orange highlights mean genes are more common in isolates from meat than human infections, green highlights mean more common in isolates from human infections than meat.

## Discussion

To our knowledge, this is the first study to phylogenetically compare ABR *E. coli* from retail meats sold for consumption by human and canine populations within a confined geographical location with contemporaneously isolated locally sourced clinical isolates.

Of meat sold at large chain grocery stores, chicken had the highest positivity for ABR *E. coli*, particularly for FQs, which, despite methodological differences between studies, is consistent with the literature (10,12,24-26). For RDF, our findings align with a recent report in the UK (27). Other international studies report similar trends, although positivity rates for 3GC-R *E. coli* vary by region (13,14,28). RDF is commonly sold frozen, and many owners incorrectly believe that freezing eliminates pathogens (7). We therefore add to existing evidence that ABR *E. coli* remain viable post-thawing of RDF (7,27,28).

Our phylogenetic analysis of ABR *E. coli* from chicken meat and chicken-based RDF suggested a shared source, consistent with the fact that UK RDF is derived from the same meat production chain as meat sold for human consumption (29). Genetic relatedness was also observed between RDF and beef meat, possibly due to mixed protein content in the RDF.

We have previously reported that excretion of FQ-R ST10, ST162 and ST744 *E. coli* by dogs in Bristol is associated with them being fed raw meat (13,14). The fact that we commonly found these same STs contaminating raw meat and RDF being sold in Bristol provides a potential explanation.

We identified four incidences where *E. coli* isolates from meat were within 20 SNPs of human clinical isolates, the closest being two ST69 chicken isolates with 12 and 13 SNPs different from a BSI isolate, with an ST69 beef isolate differing from a UTI isolate by only 16 SNPs. In each case both members of the meat/infection pair carried identical ARGs, further suggestive of isolate sharing. To our knowledge, these represent the closest genomic relationships of *E. coli* from meat and human extraintestinal infection isolates reported. This likely reflects the scarcity of high-resolution phylogenetic comparisons between human and meat/livestock isolates within the same region and time frame. One UK study (30) of similar scale to ours, reported the closest human infection to livestock relationship as 85 SNPs. However, the farm isolates were from an approximately 100 x 100 km region of England, the meat isolates (n=20) were from one city within that region, but the human infection isolates were sourced from across England, collected four to fourteen years prior to the farm/meat isolates (30). Our findings, therefore, based on a contemporaneously collected sample of meat/RDF (n=200) and human clinical isolates from one city (approximately 10 x 10 km) provide insights into potential farm-to-fork transmission routes and highlight the importance of locally integrated, high-resolution One Health genomic surveillance for their investigation.

Limitations of our study include the cross-sectional design, which cannot establish causality or transmission direction, and the relatively small sample size of meat isolates, which may underrepresent the full diversity of *E. coli* in these sources. Our study also does not capture the commensal *E. coli* present in the gastrointestinal tracts of healthy humans.

Nonetheless, our findings have important implications for public health and food safety policy. First, they underscore the potential benefits of tighter microbiological standards and regulatory oversight of RDF, which, unlike meat sold for human consumption, is explicitly sold to be fed uncooked. Second, they support including RDF in ABR bacterial surveillance as part of an integrated One Health approach. Third, public health messaging should better inform pet owners of the risks of raw feeding, and about hygiene precautions to limit household transmission.

## Transparency declaration

M.B.A. is married to the owner of a veterinary practice that sells various mass-manufactured dog foods amounting to a value less than 5% of total turnover. Otherwise, the authors declare no competing interests.

## Author Contributions

Conceived the Study: J.E.S., M.B.A.; Funding Acquisition: P.W., M.B.A.; Collection of Data: J.E.S., B.A., K.L.S., A.M.D, L.W., D.K., P.W.; Cleaning and Analysis of Data: J.E.S., M.B.A.; Initial Drafting of Manuscript: J.E.S., M.B.A.; Corrected and Approved Manuscript: All authors

Orange highlights mean genes are more common in isolates from meat than human infections, green highlights mean more common in isolates from human infections than meat.

