## Supplementary Data for "Similarities Between Antibiotic-Resistant *Escherichia coli* from Raw Meat, Commercial Raw Dog Food and Those Causing Extraintestinal Human Infections: A Contemporaneous Geographically Focussed Genomic Epidemiology Study"

**Table S1.** Reference sequences used for phylogenetic alignment of *E. coli*

| ST | ***E. coli* Reference Genome Biosample** |
|---|---|
| 10 | SAMN12231000 |
| 38 | SAMN08498861 |
| 40 | SAMN12275742 |
| 58 | SAMN12214764* |
| 69 | SAMN40242201 |
| 88 | SAMN08094773 |
| 93 | SAMN13714347 |
| 101 | SAMN10225034 |
| 117 | SAMN08993917 |
| 131 | SAMEA2272019 |
| 135 | SAMEA3138414 |
| 155 | SAMN12214764* |
| 162 | SAMN12384979 |
| 226 | SAMN26309432 |
| 349 | SAMN07618121 |
| 362 | SAMN22968086 |
| 450 | SAMN13423059 |
| 641 | SAMN13242665 |
| 1670 | SAMEA6657438 |
| 2165 | SAMN13012380 |

*ST58 and ST155 part of the same clonal complex

**Table S2**. Samples of meat sold at large chain grocery stores positive for *E. coli* on TBX without antibiotics and with the six test antibiotics.

| | | **Positive for *E. coli* selected on** | | | | | |
|---|---|---|---|---|---|---|---|
| **Sample ID** | **TBX only** | **AMX** | **SPC** | **STR** | **AMC** | **CTX** | **CIP** |
| B1.A | + | + | + | + | + | + | - |
| B1.B | + | + | + | + | - | - | - |
| B1.C | + | + | - | + | - | - | + |
| B2.A | + | - | - | - | - | - | - |
| B2.B | + | + | - | + | - | - | + |
| B2.C | + | + | + | - | + | + | - |
| B3.A | - | - | - | - | - | - | - |
| B3.B | - | - | - | - | - | - | - |
| B3.C | + | - | + | + | - | - | - |
| B4.A | + | - | - | + | - | - | - |
| B4.B | + | - | - | + | - | - | - |
| B4.C | - | - | - | - | - | - | - |
| B5.A | + | - | - | - | - | - | - |
| B5.B | + | - | - | - | - | - | - |
| B5.C | + | - | - | - | - | - | - |
| C1.A | + | + | + | + | + | - | - |
| C1.B | + | + | + | + | + | - | + |
| C1.C | + | + | + | + | + | - | + |
| C2.A | + | + | + | + | + | - | + |
| C2.B | + | + | + | + | + | - | - |
| C2.C | + | + | + | + | + | + | + |
| C3.A | + | + | + | + | + | - | - |
| C3.B | + | + | + | + | + | - | + |
| C3.C | + | + | + | + | + | - | + |
| C4.A | + | + | + | + | + | - | - |
| C4.B | + | + | + | + | + | - | - |
| C4.C | + | + | + | + | - | - | - |
| C5.A | + | + | + | + | - | - | + |
| C5.B | + | - | + | + | - | - | - |
| C5.C | + | + | + | + | + | - | - |
| L1.A | + | + | - | + | + | - | - |
| L1.B | + | + | + | + | + | - | - |
| L1.C | + | - | - | - | - | - | - |
| L2.A | + | - | + | + | - | - | - |
| L2.B | - | - | - | - | - | - | - |
| L2.C | + | + | + | + | + | - | + |
| L3.A | - | - | - | - | - | - | - |
| L3.B | + | + | - | + | - | - | - |

| | | | | | | | |
|---|---|---|---|---|---|---|---|
| L3.C | + | - | - | - | - | - | - |
| L4.A | + | - | - | - | - | - | - |
| L4.B | - | - | - | - | - | - | - |
| L4.C | + | + | + | + | - | - | - |
| L5.A | - | - | - | - | - | - | - |
| L5.B | + | - | - | - | - | - | - |
| L5.C | + | - | - | - | - | - | - |
| P1.C | + | + | + | - | + | - | - |
| P2.A | + | + | + | + | - | - | - |
| P2.B | - | - | - | + | - | - | - |
| P2.C | + | + | + | + | + | - | + |
| P3.A | - | - | - | - | - | - | - |
| P3.B | - | - | - | - | - | - | - |
| P3.C | - | - | - | - | - | - | - |
| P4.A | + | - | - | + | - | - | - |
| P4.B | + | + | + | - | - | - | - |
| P4.C | + | - | - | - | - | - | - |
| P5.A | + | + | + | + | - | - | - |
| P5.B | + | - | - | - | - | - | - |
| P5.C | + | - | - | - | - | - | - |

AMX, Amoxicillin; SPC, Spectinomycin; STR, Streptomycin; AMC, Amoxicillin-Clavulanate; CTX, Cefotaxime; CIP, Ciprofloxacin.

B, Beef; C, Chicken; L, Lamb; P, Pork. Postcode region is designated with a number and each store within that region is designated A-C.

**Table S3**. Samples of chicken-based RDF positive for *E. coli* on TBX without antibiotics and with the six test antibiotics.

| | Positive for *E. coli* in | | | | | | |
|---|---|---|---|---|---|---|---|
| **Sample ID** | **TBX** | **AMX** | **SPC** | **STR** | **AMC** | **CTX** | **CIP** |
| RDF1* | + | + | + | + | - | + | + |
| RDF2# | + | - | + | + | - | - | - |
| RDF3 | + | + | + | + | - | - | - |
| RDF4 | + | + | + | + | - | - | + |
| RDF5^ | - | - | - | - | - | - | - |
| RDF6* | + | + | + | + | + | + | + |
| RDF7# | + | + | + | + | - | - | - |
| RDF8 | - | - | - | - | - | - | - |
| RDF9 | + | + | + | + | + | + | + |
| RDF10 | + | + | + | + | - | - | + |
| RDF11* | + | + | + | + | + | + | + |
| RDF12 | + | + | + | + | - | - | - |
| RDF13 | + | + | + | + | - | - | - |
| RDF14^ | + | + | - | + | - | - | - |
| RDF15 | + | + | + | + | + | - | + |
| Positivity rate (%) | 13<br>87% | 12<br>80% | 12<br>80% | 13<br>87% | 4<br>27% | 4<br>27% | 7<br>47% |

AMX, Amoxicillin; SPC, Spectinomycin; STR, Streptomycin; AMC, Amoxicillin-Clavulanate; CTX, Cefotaxime; CIP, Ciprofloxacin.

Sample ID with symbols *, #, ^ denote multiple instances of the same brand of RDF purchased at different stores.

**Table S4**. ABR *E. coli* sequence types identified in sources of raw meat. ST found in more than one meat source in bold.

| Meat sample | Total no. of isolates | No. of STs | STs identified from individual meat source |
|---|---|---|---|
| RDF | 90 | 43 | 10x**ST10**, 8x**ST162**, 7x**ST69**, 6x**ST58**, 6x**ST665**, 6x**ST744**, 4x**ST155**, **3xST540**, 2x**ST101**, 2xST641, 2xST1140, 2xST7529, 2xST14244, 1xST; **57**, 93, **117**, 135, 224, 226, 349, **362**, 446, 450, 484, **602**, **752**, **770**, 947, 1072, 1139, 1148, 1494, 1594, 1670, 1711, **1727**, 2509, 2673, 2739, **2792**, 3941, 4616, 5662. |
| Chicken | 66 | 30 | 9x**ST10**, 7x**ST69**, 7x**ST752**, 6x**ST155**, 4x**ST665**, 3x**ST101**, 3x**ST57**, 2xST38, 2x**ST770**, 2xST1084, 2xST2165, 1xST; 40, **58**, **162**, 212, 219, 354, **362**, **602**, 1011, 1163, 1286, 1564, 1656, 2705, **2792**, 3258, 5229, 5451, 6115. |
| Beef | 17 | 11 | 5x**ST10**, 2x**ST69**, 2x**ST162**, 1xST: 109, **131***, 154, **540**, 906, 1204, 1252, **1727**. |
| Lamb | 13 | 12 | 2xST201, 1xST: 43, **88**, 111, **117**, 156, **162**, 1326, **1718**, 2073, 2179, novel 5597. |
| Pork | 14 | 11 | 3x**ST88** ,2x**ST10**, 1xST: 56, **131****, 165, 218, 542, **744**, **1718**, 5271, and an unknown ST. |

** O16:H5 fimH41 ** O15:H4 fimH22*

**Table S5.** Clones seen in meat isolates where members are <20 SNPs from at least one other member

| | | | | | | |
|---|---|---|---|---|---|---|
| **Clone 1**<br>**ST10** | 236981_B1A<br>*aadA22, tetA* | 288809_B1A<br>*aadA22, tetA, sul1, fosA7* | | | | |
| **Clone 2**<br>**ST10** | 236997_C3A<br>*aadA1, aadA2, sul1, sul3, tetA, cmlA1, floR, mefB* | 245208_C5B<br>*aadA1, aadA2b, sul1, dfrA1, tetA, inuF* | 246473_C4A<br>*aadA1, sul2, dfrA1, blaTEM-1* | | | |
| **Clone 3**<br>**ST10** | 246463_B2C<br>*aadA1, blaCTX-M-15, blaOXA-1, tetB. catA1* | 246491_C5A<br>*dfrA14, blaTEM-1, tetA, floR, qnrS1* | 251333_RDF1<br>*aac(3)-IId, aadA1, aadA2, aph(3'')-Ia, strAB, sul2, sul3, dfrA12, blaTEM-1, tetB, tetM, cmlA1, floR* | | | |
| **Clone 4**<br>**ST10** | 251336_RDF1<br>*aadA5, aadA24, strAB, sul1, sul2, dfrA17, blaCTX-M-1, blaTEM-1, tetA, tetB, catA1, inuG* | 251361_RDF6<br>*aadA5, aadA24, strAB, sul1, sul2, dfrA17, blaCTX-M-1, blaTEM-1, tetA, tetB, catA1, inuG* | | | | |
| **Clone 5**<br>**ST10** | 251379_RDF9<br>*aadA12, blaTEM-1, sul1* | 254066_RDF15<br>*aadA24, blaTEM-1, inuG* | 254042_RDF11<br>*strAB, dfrA14, sul2, blaTEM-1, tetA* | | | |
| **Clone 6**<br>**ST10** | 265835_C3C<br>*aadA24, blaTEM-1, inuG* | 254056_RDF12<br>*aadA1, sul2, dfrA1, blaTEM-1, tetA* | | | | |
| **Clone 7**<br>**ST10** | 265846_C4C<br>*aadA1, blaTEM-1, inuF* | 265848_C5C<br>*aadA24, blaTEM-1, inuG* | | | | |
| **Clone 8**<br>**ST38** | 245218_C2B<br>*blaTEM-1* | 265795_C1A<br>*blaTEM-1* | | | | |
| **Clone 9**<br>**ST58** | 251360_RDF6<br>*strAB, sul2, dfrA5, blaTEM-1, tetA* | 265864_RDF11<br>*strAB, dfrA5, blaTEM-1, tetA* | | | | |
| **Clone 10**<br>**ST58/ST7529** | 245221_C3A<br>*blaTEM-1* | 251351_RDF4<br>*aadA22, sul2, dfrA1, blaTEM-1, inuF* | 251335_RDF1<br>*aadA1, ant(2'')-Ia, sul1, sul2, dfrA36, floR* | 251356_RDF6<br>*aadA1, ant(2'')-Ia, sul1, sul2, dfrA36, floR* | | |
| **Clone 11**<br>**ST69** | 236984_B3C<br>*aac(3)-IVa, aadA1, sul1, sul2* | 265833_C3B<br>*aac(3)-IVa, aadA1, sul1, sul2* | 265841_C4A<br>*aac(3)-IVa, aadA1, sul1, sul2* | 245206_C5A<br>*aac(3)-IVa, aadA1, sul1, sul2* | 254065_RDF15<br>*aac(3)-IVa, aadA1, sul1, sul2, inuG, blaTEM-1* | 254067_RDF15<br>*aac(3)-IVa, aadA1, sul1, sul2* |

| | | | | | | | | | |
|---|---|---|---|---|---|---|---|---|---|
| **Clone 12**<br>**ST69** | 236994_C2A<br>*aac(3)-Iva, aadA1, sul1, sul2* | 265827_C2C<br>*aac(3)-Iva, aadA1, sul1, sul2* | | | | | | | |
| **Clone 13**<br>**ST69** | 251368_RDF7<br>*strAB, sul2, dfrA5, blaTEM-1* | 251370_RDF7<br>*strAB, aadA2b, qnrS1, sul2, dfrA5, blaTEM-1, blaCTX-M-1, inuF, mphA* | | | | | | | |
| **Clone 14**<br>**ST69/ST14244** | 251345_RDF3<br>*blaTEM-1* | 254063_RDF15<br>*blaTEM-1* | 254062_RDF14<br>*blaTEM-1* | 251369_RDF7<br>*strAB, sul2, dfrA5, blaTEM-1, tetA* | | | | | |
| **Clone 15**<br>**ST88** | 245210_P2A<br>*strAB, sul2, tetB* | 245213_P4A<br>*strAB, sul2, dfrA5, blaTEM-1, tetB* | | | | | | | |
| **Clone 16**<br>**ST101** | 236993_C1C<br>*aadA2b, aadA24, blaTEM-1, tetA, inuF, inuG* | 265845_C4C<br>*aadA1, aadA2, sul3, dfrA12, blaTEM-1, tetA, cmlA1* | 246465_C1B<br>*aadA1, sul2, dfrA1, blaTEM-1* | 254060_RDF14<br>*strAB, blaTEM-1, tetB,* | 254053_RDF12<br>*aadA2b, blaTEM-1, inuF* | | | | |
| **Clone 17**<br>**ST155** | 246472_C3C<br>*aadA1, aadA2b, sul2, dfrA1, blaTEM-1, inuF* | 265819_C2C<br>*sul2, dfrA1, dfrA5, blaTEM-1, tetA* | 265820_C2C<br>*aadA1, sul2, dfrA1, blaTEM-1* | 265801_C1B<br>*aadA1, dfrA1, blaTEM-1, tetA, qnrS1* | 254055_RDF12<br>*aadA1, dfrA1, blaTEM-1, tetA, qnrS1* | | | | |
| **Clone 18**<br>**ST155** | 265828_C2C<br>*aac(3)-Iva, aadA1, strAB, sul1, sul2, dfrA1, blaTEM-1, tetA* | 265858_RDF6<br>*ampC -42, tetA* | | | | | | | |
| **Clone 19**<br>**ST162** | 237038_B1C<br>*strAB, sul2, tetA, floR* | 246484_B2B<br>*strAB, sul2, tetA, floR* | 246485_C1B<br>*strAB, sul2, tetA, floR* | 251364_RDF6<br>*aac(3)-Iid, aadA17, blaTEM-1, tetB, inuF* | 254041_RDF10<br>*blaTEM-1* | 251340_RDF1<br>*aadA5, aph(3'')-Ia, strAB, sul2, dfrA17, blaTEM-1, tetB* | 251366_RDF6<br>*aadA5, aph(3'')-Ia, strAB, sul2, dfrA17, blaTEM-1, tetB* | 254070_RDF15<br>*dfrA17, blaTEM-1, tetB* | 265855_RDF6<br>*blaTEM-1, tetB, catA1* |
| **Clone 20**<br>**ST201** | 246476_L1A<br>*strAB, blaTEM-1, tetB* | 288820_L1A<br>*strAB, blaTEM-1* | | | | | | | |
| **Clone 21**<br>**ST540** | 246494_B1A<br>*strAB, sul2, dfrA14, blaCTX-M-15, blaTEM-1, tetA, qnrS1* | 251354_RDF6<br>*blaTEM-1* | 265863_RDF11<br>*strAB, dfrA5, blaTEM-1, tetA, tetB* | 265860_RDF9<br>*aadA1, strAB, sul1, sul2, dfrA17, dfrA36, blaDHA-1, blaTEM-1, blaOXA-1, tetB, qnrB4, mphA* | | | | | |

| | | | | | | | | |
|---|---|---|---|---|---|---|---|---|
| **Clone 22 ST665** | 246464_C1A *blaTEM-1* | 265816_C1C *aadA22, blaTEM-1, inuF* | 265831_C3A *aadA1, strAB, sul1, sul2, dfrA1, blaTEM-1, tetA, inuF, mphB* | 265798_C1A *aadA1* | 254064_RDF15 *aadA1, strAB, sul1, sul2, dfrA1, blaTEM-1* | 265867_RDF15 *aadA1, strAB, sul1, sul2, dfrA1, blaTEM-1, tetA, qnrS1* | 265853_RDF6 *aadA1, aadA2b, aph(3')-Ia, sul2, blaTEM-1, qnrB19, inuF, mphA* | 265854_RDF6 *aadA1, aph(3')-Ia, blaTEM-1, mphA* |
| **Clone 23 ST665** | 251344_RDF3 *aadA1, aadA2b, sul3, dfrA16, blaCARB-2, cmlA1* | 254035_RDF10 *aadA1, blaTEM-1, inuF* | | | | | | |
| **Clone 24 ST744** | 246493_P2C *aadA12, aph(3")-Ib, strAB, sul1, sul2, blaTEM-1, tetA, tetB, mphA* | 251341_RDF1 *strAB, sul2, tetB, catA1* | 251363_RDF6 *strAB, sul2, tetB, catA1* | 254049_RDF11 *aadA5, strAB, sul1, sul2, dfrA17, blaTEM-1, tetA, catA1, floR, mphA* | 254050_RDF11 *aadA5, strAB, sul1, sul2, dfrA17, blaTEM-1, tetB, catA1, mphA* | 251383_RDF9 *aadA5, aph(3")-Ib, strAB, sul1, sul2, dfrA17, blaTEM-1, tetB, catA1, mphA* | 251380_RDF9 *aadA5, aph(3")-Ib, strAB, sul1, sul2, dfrA17, blaTEM-1, tetA, catA1, floR, mphA* | |
| **Clone 25 ST752** | 236996_C2C *aadA1, strAB, sul1, blaTEM-1, inuF* | 265837_C3C *aadA1, strAB, sul1, blaTEM-1, inuF* | 245209_C5C *aadA1, strAB, sul1, blaTEM-1* | 265839_C4A *aadA2b, strAB, blaTEM-1, inuF* | 245223_C4C *strAB, blaTEM-1, qnrB, qnrS1* | 265840_C4A *aadA12, strAB, blaTEM-1* | 265847_C4C *strAB, sul2, blaTEM-1* | 251365_RDF6 *strAB* |
| **Clone 26 ST770** | 246470_C3A *aac(3)-IVa, aadA1, sul1, sul2, blaTEM-1* | 251362_RDF6 *aadA1, aadA2, aadA17, sul2, sul3, dfrA12, blaSHV-12, tetA, cmlA1* | | | | | | |
| **Clone 27 ST1084** | 236992_C1B *aac(3)-IVa, aadA1, strAB, sul1, blaTEM-1, tetA, cmlA1, mcr10* | 265806_C1B *aac(3)-IVa, aadA1, sul1, tetA* | | | | | | |
| **Clone 28 ST1727** | 246462_B1A *fosA7* | 251367_RDF7 *aadA1, sul1, sul2, dfrA36, blaOXA-1, tetB, floR, fosA7* | | | | | | |
| **Clone 29 ST2165** | 265823_C2C *aadA1, aadA17, aadA2b, sul3, blaSHV-12, cmlA, tetA* | 265824_C2C *aadA1, aadA5, aadA2b, dfrA17, sul2, sul3, blaSHV-12, cmlA, tetA* | | | | | | |
| **Clone 30 ST2792** | 265807_C1B *aadA12, blaTEM-1* | 254057_RDF13 *aadA12, blaTEM-1* | | | | | | |

Blue: Beef, "B", Light orange: Chicken, "C", Dark orange: "RDF", Pink: Pork, "P", Grey: Lamb, "L". Sample numbers are 1A-C to 5A-C for grocery store meat and 1 to 15 for RDF

**Table S6.** STs (number of isolates) shared between *E. coli* from human BSI/UTI and at least one type of meat

| HUMAN | RDF | Chicken | Red Meat |
|---|---|---|---|
| 10 | 10 (10) | 10 (9) | 10 (7) |
| 38 | | 38 (2) | |
| 40 | | 40 (1) | |
| 58 | 58 (6) | 58 (1) | |
| 69 | 69 (7) | 69 (7) | 69 (2) |
| 88 | | | 88 (4) |
| 93 | 93 (1) | | |
| 101 | 101 (2) | 101 (3) | |
| 117 | 117 (1) | | 117 (1) |
| 131 | | | 131 (2) |
| 135 | 135 (1) | | |
| 155 | 155 (4) | 155 (6) | |
| 162 | 162 (8) | 162 (1) | 162 (3) |
| 226 | 226 (1) | | |
| 349 | 349 (1) | | |
| 362 | 362 (1) | 362 (1) | |
| 450 | 450 (1) | | |
| 542 | | | 542 (1) |
| 641 | 641 (2) | | |
| 1670 | 1670 (1) | | |
| 2165 | | 2165 (2) | |

**Table S7.** Number of *E. coli* phylogroups identified from poultry and red meat and from human BSI and UTI from the same city collected in parallel.

| **Phylogroup** | **Poultry** | | **Red Meat** | | **Human** | |
|---|---|---|---|---|---|---|
| A/B1 | 103 | 66.0% | 30 | 68.2% | 159 | 13.5% |
| B2 | 1 | 0.61% | 3 | 6.82% | 793 | 67.1% |
| C | 1 | 0.61% | 5 | 11.4% | 12 | 1.02% |
| D | 35 | 22.4% | 3 | 6.82% | 163 | 13.8% |
| E | 12 | 7.69% | 1 | 2.27% | 0 | 0.00% |
| G/F | 4 | 2.56% | 1 | 2.27% | 43 | 3.64% |
| UNKNOWN | 0 | 0.00% | 1 | 2.27% | 12 | 1.02% |

**Table S8.** Number of incidences where *E. coli* from meat are <=50 SNPs from *E. coli* causing human infection (=no. of isolates)

| Clone | Human | Meat | SNPs | ST |
|---|---|---|---|---|
| 1* | 254113_23M70147151 | 251368_RDF72 | 48 | ST69 |
| 2* | 267852_23M00132532 | 254036_RDF102 | 37 | ST69 |
| | 272261_23M80061343 | 254036_RDF102 | 41 | ST69 |
| 3* | 273974_23M60199005 | 236994_C2Aspec | 12 | ST69 |
| | 273974_23M60199005 | 265827_C2CT9 | 13 | ST69 |
| 4* | 266036_US534 | 245214_B4Bstrep | 16 | ST69 |
| 5 | 255797_US18 | 246478_L2CAMC | 30 | ST88 |
| 6 | 254166_23M00030325 | 254059_RDF133 | 36 | ST117 |
| | 255610_23M70185683 | 254059_RDF133 | 45 | ST117 |
| | 255809_US30 | 254059_RDF133 | 22 | ST117 |
| | 272048_23M80044177 | 254059_RDF133 | 19 | ST117 |
| | 274038_23M00315677 | 254059_RDF133 | 44 | ST117 |
| 7 | 249681_23M80027438B | 246483_B1CCIP | 49 | ST131 |
| 8** | 258393_US513 | 251340_RDF19 | 40 | ST162 |
| | 258393_US513 | 251366_RDF614 | 41 | ST162 |
| | 258393_US513 | 254070_RDF158 | 44 | ST162 |
| | 272106_23M00189152 | 251340_RDF19 | 45 | ST162 |
| | 272106_23M00189152 | 251366_RDF614 | 44 | ST162 |
| | 272106_23M00189152 | 254070_RDF158 | 45 | ST162 |

*Isolates from clone 1 to clone 4 are 88 – 385 SNPs apart

**RDF Isolates are part of the extensive clone 19 of meat isolates with <20 SNPs

**Table S9**. Resistance genes found in ABR *E. coli* from sources of raw meat

| Resistance Genes | RDF | Chicken | Beef | Pork | Lamb |
|---|---|---|---|---|---|
| Beta-lactams | | | | | |
| *blaCARB-2* | 4 | 1 | 0 | 0 | 0 |
| *blaCMY-100* | 0 | 0 | 0 | 1 | 0 |
| *blaCTX-M-1* | 4 | 0 | 0 | 0 | 0 |
| *blaCTX-M-15* | 3 | 0 | 3 | 0 | 0 |
| *blaDHA-1* | 1 | 0 | 0 | 0 | 0 |
| *blaSHV-12* | 2 | 2 | 0 | 0 | 0 |
| *blaTEM-1A* | 4 | 1 | 2 | 1 | 0 |
| *blaTEM-1B* | 56 | 48 | 7 | 2 | 8 |
| *blaTEM-1C* | 1 | 1 | 0 | 2 | 1 |
| *blaTEM-1D* | 1 | 2 | 0 | 0 | 0 |
| *blaTEM-78* | 1 | 0 | 0 | 0 | 0 |
| *blaTEM-106* | 0 | 1 | 1 | 1 | 0 |
| *blaTEM-120* | 0 | 1 | 0 | 0 | 0 |
| *blaTEM-126* | 0 | 1 | 1 | 1 | 0 |
| *blaTEM-135* | 0 | 1 | 1 | 1 | 0 |
| *blaTEM-176* | 0 | 0 | 0 | 1 | 0 |
| *blaTEM-190* | 1 | 0 | 0 | 0 | 0 |
| *blaTEM-220* | 0 | 0 | 1 | 1 | 0 |
| *blaOXA-1* | 2 | 0 | 1 | 0 | 0 |
| Fluoroquinolones | | | | | |
| *qnrB4* | 1 | 0 | 0 | 0 | 0 |
| *qnrB5* | 0 | 1 | 0 | 0 | 0 |
| *qnrB19* | 1 | 3 | 0 | 0 | 0 |
| *qnrB21* | 0 | 0 | 0 | 1 | 0 |
| *qnrB81* | 0 | 1 | 0 | 0 | 0 |
| *qnrS1* | 7 | 3 | 1 | 1 | 0 |
| Gentamicin/Tobramycin | | | | | |
| *aac(3)-IId* | 3 | 0 | 0 | 0 | 1 |
| *aac(3)-IV* | 0 | 1 | 1 | 0 | 0 |
| *aac(3)-VIa* | 2 | 10 | 0 | 0 | 0 |
| *ant(2")-Ia* | 3 | 0 | 0 | 0 | 0 |
| Trimethoprim | | | | | |
| *dfrA1* | 12 | 15 | 1 | 3 | 1 |
| *dfrA5* | 8 | 1 | 0 | 1 | 1 |
| *dfrA7* | 1 | 0 | 1 | 0 | 0 |
| *dfrA12* | 4 | 2 | 0 | 0 | 0 |
| *dfrA14* | 3 | 2 | 1 | 1 | 0 |
| *dfrA16* | 2 | 1 | 0 | 0 | 0 |
| *dfrA17* | 12 | 1 | 1 | 0 | 1 |
| *dfrA36* | 7 | 0 | 0 | 0 | 0 |
| *dfrB1* | 0 | 0 | 0 | 1 | 0 |
| Sulphonamides | | | | | |
| *sul1* | 25 | 20 | 6 | 5 | 2 |
| *sul2* | 48 | 25 | 9 | 5 | 3 |
| *sul3* | 6 | 6 | 0 | 0 | 0 |
| Tetracycline | | | | | |
| *tet(A)* | 36 | 21 | 13 | 7 | 4 |
| *tet(B)* | 25 | 4 | 2 | 4 | 2 |
| *tet(C)* | 0 | 0 | 0 | 1 | 0 |

| | | | | | |
|---|---|---|---|---|---|
| *tet(M)* | 2 | 0 | 0 | 0 | 0 |
| Florfenicol | | | | | |
| *catA1* | 12 | 1 | 2 | 0 | 0 |
| *cmlA1* | 6 | 7 | 0 | 0 | 0 |
| *floR* | 13 | 2 | 3 | 1 | 0 |
| Streptomycin/Spectinomycin/Neomycin | | | | | |
| *aadA1* | 30 | 38 | 4 | 6 | 2 |
| *aadA2* | 4 | 5 | 0 | 0 | 0 |
| *aadA2b* | 7 | 7 | 0 | 0 | 0 |
| *aadA5* | 10 | 1 | 1 | 0 | 0 |
| *aadA12* | 2 | 3 | 0 | 1 | 0 |
| *aadA17* | 4 | 1 | 0 | 0 | 0 |
| *aadA22* | 3 | 2 | 3 | 0 | 3 |
| *aadA23* | 0 | 0 | 0 | 1 | 0 |
| *aadA24* | 5 | 4 | 0 | 0 | 1 |
| *ant(3")-Ia* | 0 | 2 | 0 | 0 | 0 |
| *aph(3")-Ia* | 8 | 0 | 0 | 3 | 1 |
| *aph(3")-Ib* | 38 | 13 | 9 | 5 | 6 |
| *aph(4)-Ia* | 0 | 1 | 0 | 0 | 0 |
| *aph(6)-Id* | 38 | 16 | 9 | 5 | 6 |
| Others | | | | | |
| *lnu(F)* | 8 | 17 | 0 | 0 | 1 |
| *lnu(G)* | 8 | 8 | 0 | 0 | 1 |
| *ere(B)* | 2 | 0 | 0 | 0 | 0 |
| *erm(B)* | 0 | 0 | 1 | 0 | 0 |
| *mph(A)* | 10 | 0 | 1 | 1 | 0 |
| *mph(B)* | 0 | 2 | 1 | 1 | 0 |
| *mef(B)* | 0 | 1 | 0 | 0 | 0 |
| *mph(E)* | 1 | 0 | 0 | 0 | 0 |
| *msr(E)* | 1 | 0 | 0 | 0 | 0 |
| *fosA7* | 1 | 2 | 2 | 0 | 0 |
| *mcr-10* | 0 | 1 | 0 | 0 | 0 |
| Total | 499 | 310 | 89 | 65 | 45 |
| Avg no. R genes/ isolate | 5.54 | 4.70 | 5.24 | 5.00 | 3.21 |
| Mutations | | | | | |
| *ampC* -42 | 1 | 0 | 0 | 0 | 0 |
| *gyrA*p.D87N | 18 | 6 | 3 | 1 | 0 |
| *gyrA*p.D87Y | 1 | 0 | 0 | 0 | 1 |
| *gyrA*p.S83A | 1 | 4 | 0 | 0 | 1 |
| *gyrA*p.S83L | 24 | 13 | 3 | 1 | 0 |
| *parC*p.A56T | 6 | 0 | 0 | 1 | 0 |
| *parC*p.E84G | 1 | 1 | 0 | 0 | 0 |
| *parC*p.E84K | 0 | 1 | 0 | 0 | 0 |
| *parC*p.E84V | 0 | 0 | 1 | 1 | 0 |
| *parC*p.S57T | 2 | 3 | 0 | 0 | 0 |
| *parC*p.S80I | 20 | 6 | 3 | 1 | 1 |
| *parC*p.S80R | 0 | 1 | 0 | 0 | 0 |

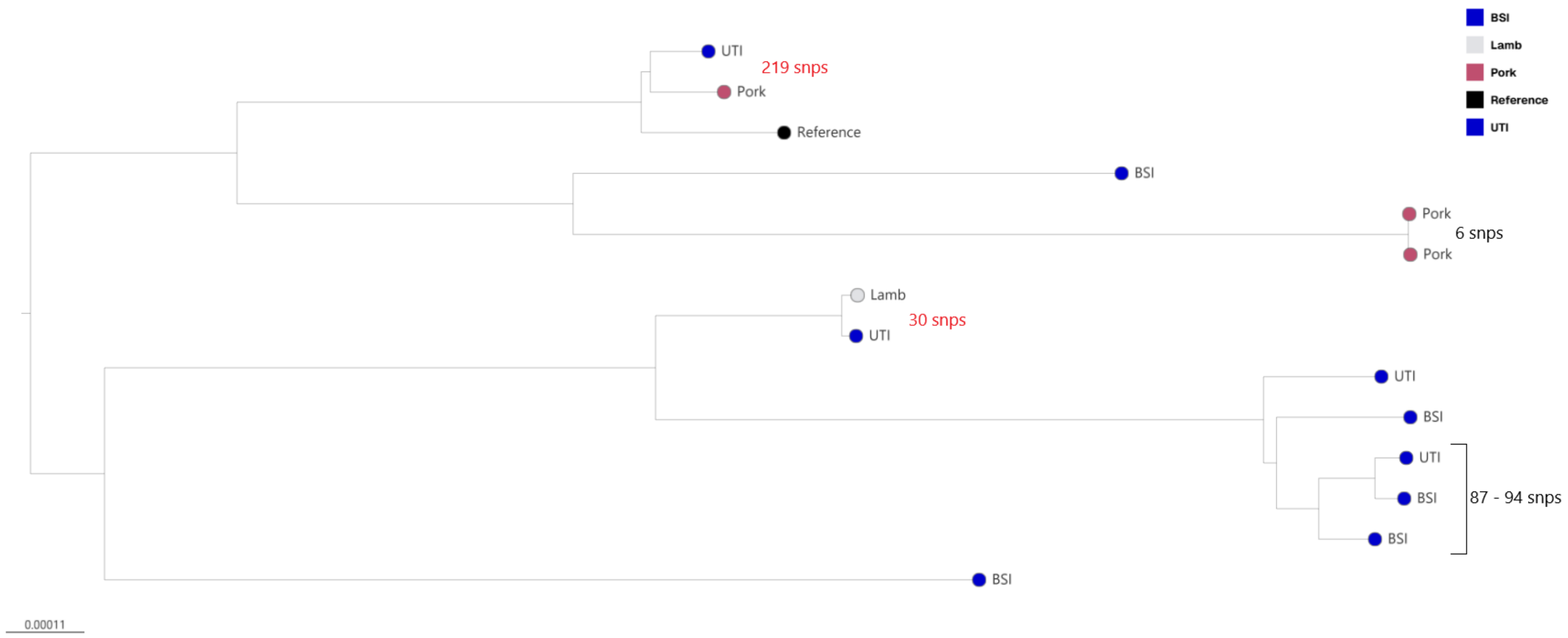


**Figure S1.** Mid-rooted phylogenetic tree of *E. coli* ST88 isolates from human clinical infections (HCI; bloodstream and urinary tract infections, blue), pork (pink) and one lamb (grey). Single nucleotide polymorphisms in red are SNPs between raw meat source and human source, SNPs between isolates from one source in black.

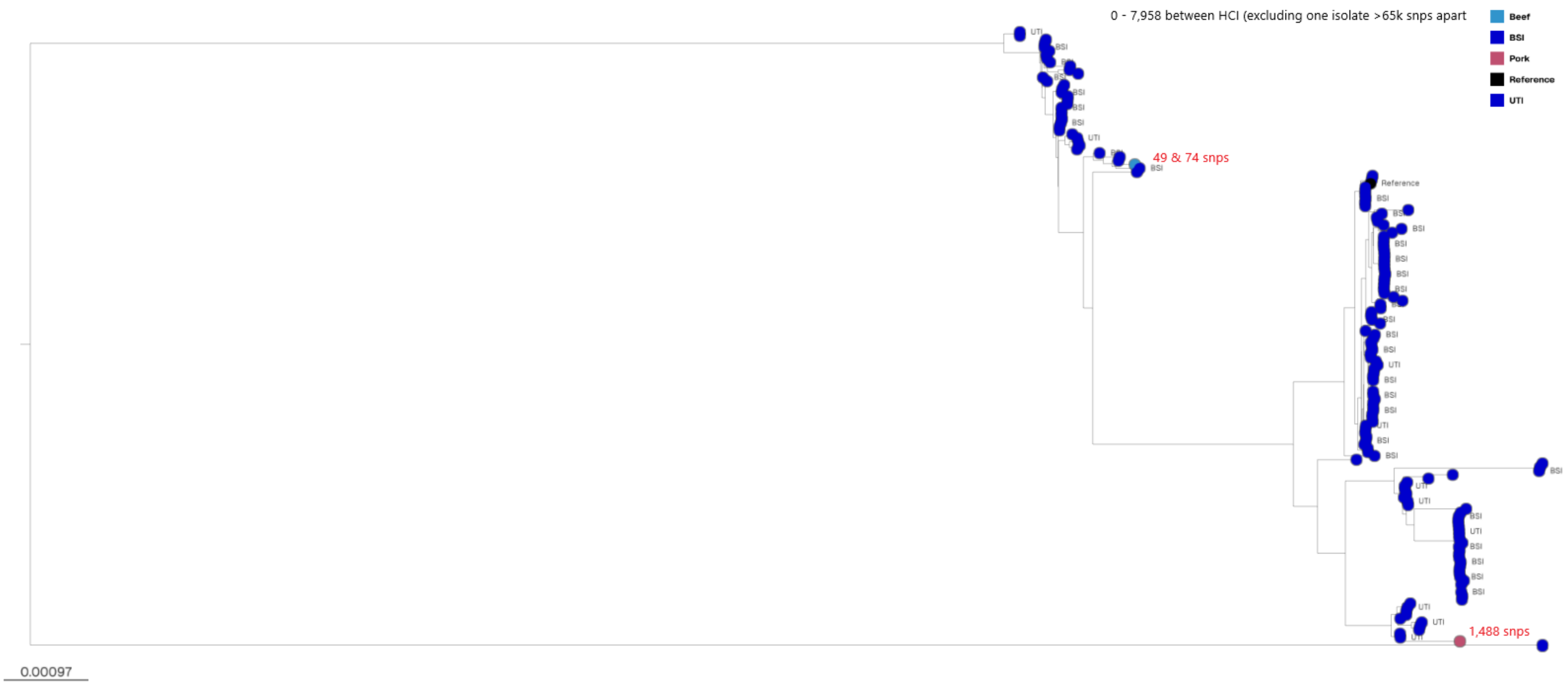


**Figure S2.** Mid-rooted phylogenetic tree of *E. coli* ST131 isolates from human clinical infections (HCI; bloodstream and urinary tract infections, blue), and a beef (blue) and pork isolate (pink). Closest single nucleotide polymorphisms in red are SNPs between raw meat source and human source, SNPs between HCI isolates in black.

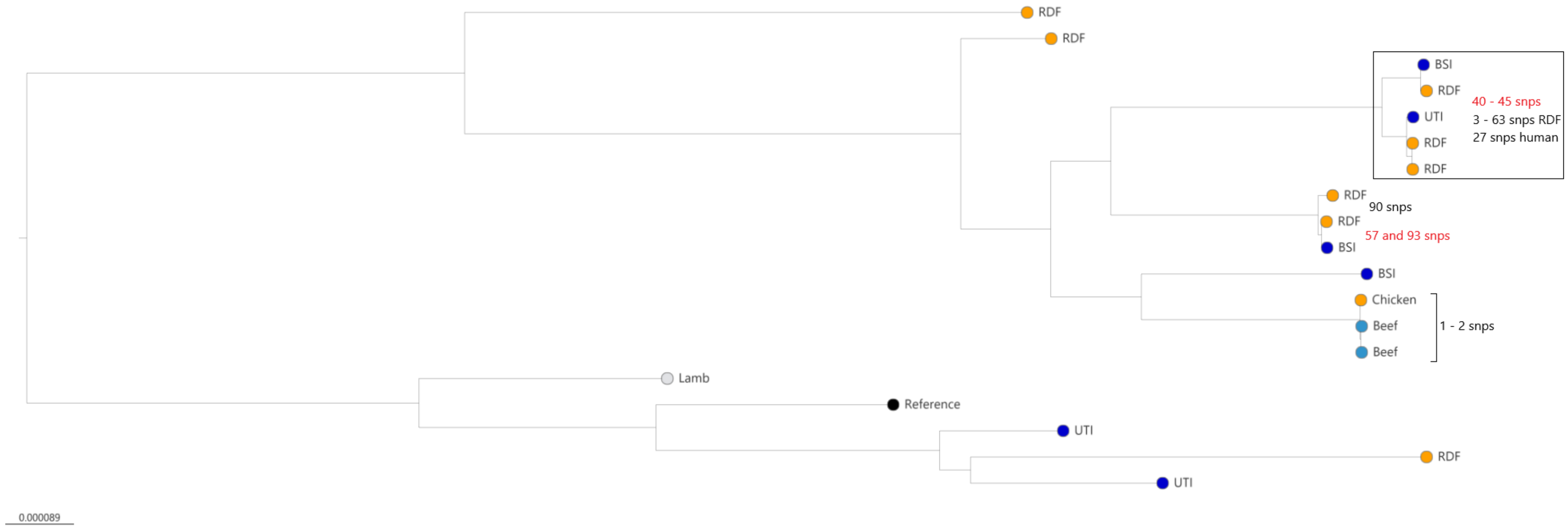


**Figure S3** Mid-rooted phylogenetic tree of *E. coli* ST162 isolates from human bloodstream and urinary tract infections, and from raw meat sources. RDF= raw dog food. Single nucleotide polymorphisms in red are SNPs between raw meat source and human source, SNPs between isolates from one source in black.
